# Human microbiota is a reservoir of SARS-CoV-2 advantageous mutations

**DOI:** 10.1101/2023.11.16.567485

**Authors:** Birong Cao, Xiaoxi Wang, Wanchao Yin, Zhaobing Gao, Bingqing Xia

## Abstract

SARS-CoV-2 mutations are rapidly emerging, in particular advantageous mutations in the spike (S) protein, which either increase transmissibility or lead to immune escape, are posing a major challenge to pandemic prevention and treatment. However, how the virus acquires a high number of advantageous mutations in a short time remains a mystery. Here, we show that the human microbiota may contribute to mutations in variants of concern (VOCs). We identified a mutation and adjacent 6 amino acids (aa) in a viral mutation fragment (VMF) and searched for homologous fragments (HFs) in the National Center for Biotechnology Information (NCBI) database. Among the approximate 8000 HFs obtained, 61 mutations in S and other outer membrane proteins were found in bacteria, accounting for 62% of all mutation sources, which is a 12-fold higher than the natural variable proportion. Approximately 70% of these bacterial species belong to the human microbiota, are primarily distributed in the gut or lung and exhibit a composition pattern similar to that of COVID-19 patients. Importantly, SARS-CoV-2 RNA-dependent RNA polymerase (RdRp) replicates corresponding bacterial mRNAs harboring mutations, producing chimeric RNAs. Collectively, SARS-CoV-2 may acquire mutations from the human microbiota, resulting in alterations in the binding sites or antigenic determinants of the original virus. Our study sheds light on the evolving mutational mechanisms of SARS-CoV-2.

## Text

By October 2022, amino acid (aa) mutations in severe acute respiratory syndrome coronavirus 2 (SARS-CoV-2) proteins had accumulated to thirty thousand (https://ngdc.cncb.ac.cn)^1,2^. Among these mutations, mutations in the spike (S) protein were of particular significance because the S protein mediates viral entry into a host cell and is a target for vaccine development ^3–5^. Some S mutations, such as N501Y, D614G, and L452R, show the potential to alter the efficiency of viral entry into human cells and enhance transmissibility from 10- to 100- fold ^6–10^. Mutations E484K, F490L, G446S, K417N, L452R, N501Y and S477N, located in the receptor-binding domain (RBD) of the S protein, significantly reduce the effects of neutralizing antibodies and vaccine efficacy ^4,11,12^. These mutations, which either increase transmissibility or lead to immune escape, are thought to be advantageous mutations to SARS-CoV-2. On the basis of increased transmissibility or a change in clinical disease presentation, which is primarily caused by mutations in the S protein, 9 variants have been classified as variants of concern (VOCs) by the World Health Organization (WHO): the alpha variant (originally identified in the UK, B.1.1.7), the beta variant (originally identified in South Africa, B.1.351), the gamma variant (originally identified in Brazil, P.1), the delta variant (originally identified in India, B.1.617.2) and the omicron variant (originally identified in India, BA.1-BA.5) ^13,14^. The rapid emergence of mutations poses a major challenge to the effectiveness of vaccines and therapeutic agents and has ushered in a new area of the COVID-19 pandemic ^15,16^.

Some studies support the idea that obtaining advantageous mutations is an inevitable result of SARS-CoV-2 evolution and natural selection ^17,18^. However, the short period in which VOCs have acquired the super ability to infect the global population has been astonishing. Being an enveloped positive-sense single-stranded RNA virus, SARS-CoV-2 requires an RNA-dependent RNA polymerase (RdRp) for replication, which is inherently prone to error ^3^. However, this type of replication-associated mutation is not easy to obtain via evolutionarily advantageous mutation, and although they benefit a virus by promoting host adaptation capacity, they can be fatal to the virus itself ^6,17^. Recombination among coronaviruses or across viral species may lead to a greater number of advantageous mutations than in acquired through replication, leading to significant changes in the phenotype of SARS-CoV-2 ^19,20^. However, thus far, only a few recombination events, mostly between this variant and other subspecies of coronaviruses in the same genus, have been identified, making understanding how SARS-CoV-2 acquired a high number of advantageous mutations in a short time difficult ^18,21^.

Multiple unusual “events” lead to the mystery surrounding the mutational evolution of SARS-CoV-2. On the one hand, some VOCs share identical dominant mutations, such as N501Y and D164G in the alpha, beta, gamma and omicron variants and K417N and L452R in the delta and omicron variants, supporting the evolutionary relationships of these VOCs ^6,22^. However, some mutations arose independently in different geographical locations at the same time, such as N501Y, which emerged in England and South Africa at the same time ^23,24^. Finally, compared to other VOCs, the omicron variant appears to be unique. Omicron carries 25 unique mutations in the S protein, which suggests that some of these mutations were not derived from variants circulating before and therefore might have a different origin. The rapid accumulation of many mutations made the omicron variant an ideal virus to use for investigating mutation origins ^25,26^. One hypothesis suggests that the omicron variant may have ‘cryptically spread’ for a long time and circulated in a population with insufficient viral surveillance and sequencing, suggesting a similar evolutionary mechanism to the other VOCs ^27,28^. Another hypothesis suggests that the Omicron variant may have evolved in immunocompromised COVID-19 patients, supporting an intrahost evolution during prolonged infection ^29–31^. Finally, a third hypothesis posits that the omicron variant may have accumulated mutations in a nonhuman host because the sequence of the omicron S protein significantly overlaps with SARS-CoV-2 that carries mutations known to promote adaptation in mice ^32,33^.

Nerveless, various signs have made clear that SARS-CoV-2 rapidly acquired multiple advantageous mutations, which promoted its evolution toward increased adaptiveness, evolving almost perfectly in this direction. However, the current hypotheses and theories are insufficient to explain how SARS-CoV-2 acquired a high number of mutations, particularly those with evolutionary advantages, that has enabled the virus to spread in the face of increasing population immunity while maintaining or increasing replication fitness. Therefore, it has been speculated that currently unrecognized factors have facilitated SARS-CoV-2 evolution into a better adaptive species? If these factors account for the rapid mutation, they would be expected to meet the following characteristics: widely distributed in hosts, contribute to easy viral access to hosts, and perhaps is immunologically tolerant to hosts. The potential factors that facilitate advantageous mutation are explored.

## Results

### A viral mutation fragment-based blast was developed to identify homologous fragments harboring advantageous mutations

The prevalence of S protein mutations in all VOCs was first analyzed. We found that among the identified 3116 S protein mutations, all 55 advantageous mutations had a mutation frequency higher than 90% ^1,2^. The S protein comprises two domains: the S1 domain, which contains the RBD, and the S2 domain, which is critical for membrane fusion ^5^. The RBD mediates attachment to human cells and is the primary target of neutralizing antibodies, and it carries 90% of S mutations, including the intensively studied N501Y, E484K, L452Q and F486V (Fig. 1a) ^12,34^. In addition, some mutations in addition to those of the S protein may markedly affect the direction of the pandemic; these include T9I, a mutation in the membrane envelope (E) protein, leading to virulence reduction in the Omicron variant ^35^. This finding inspired us to include the high frequency mutations in E and membrane (M) proteins, two other structural membrane proteins, constituting the outer protein envelope and S protein together. Ultimately, a total of 61 mutations were identified: 55 S protein mutations, 2 E protein mutations and 4 M protein mutations (Fig. 1b, Fig. S1a). Notably, the omicron variants, including BA.4 and BA.5 subvariants, carry 38 mutations, which is a much higher number than the that of any other VOCs (Fig. S1, Table. S1b).

**Fig. 1.**
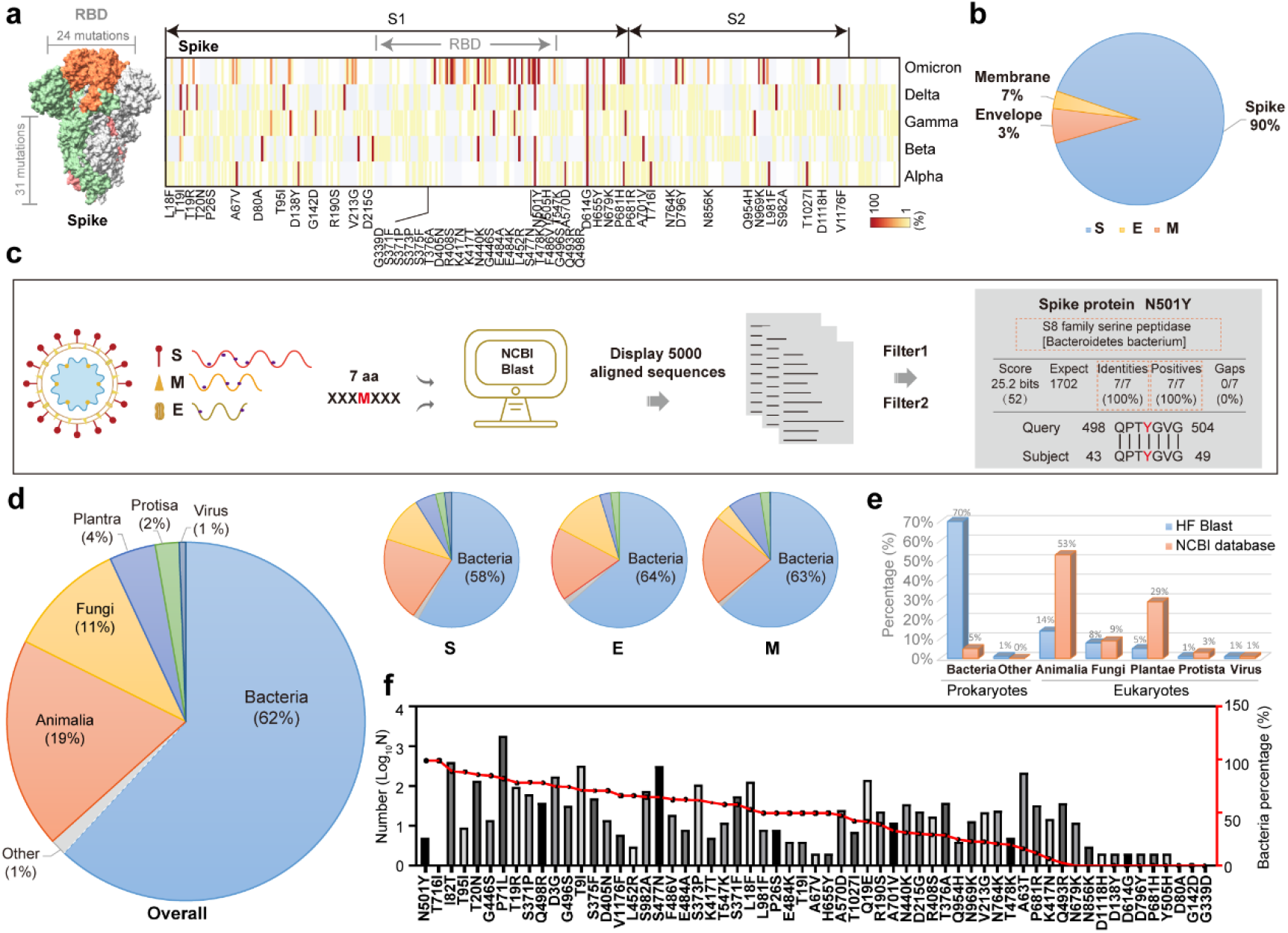
Homologous fragment (HF) alignment of viral mutation fragments (VMFs) harboring high-frequency mutations in outer membrane proteins. **a,** Structure of the SARS-CoV-2 S protein. The RBD domain containing 24 mutations is marked in orange (left). The probability of S protein mutation in all VOCs (right). **b,** The ratio of high-frequency mutations in the spike (S), envelope (E) and membrane (M) proteins. **c,** Flow chart showing homologous sequence alignment of viral mutation fragments in the NCBI database. **d,** Pie chart representations of HF kingdoms derived from overall (left bigger), spike (S), envelope (E) and membrane (M) (right smaller three) mutations. **e,** The species proportions contributing HFs and in NCBI databases. **f,** Correlation analysis of the number of HFs for each mutation and the corresponding bacterial contribution.

In contrast to focusing on a single amino acid mutation itself, we focused on the corresponding viral mutation fragment (VMF), which includes a mutated amino acid and adjacent amino acids as a unit. A previous study showed that PRRA, a VMF containing 4 amino acids (aa), functioned as a furin cleavage site ^36,37^. The length of a VMF was tentatively set to be 5 aa, and a VMF-based blast, aiming to identify homologous fragments (HFs) of these VMFs, was carried out with the NCBI Protein database, the largest protein database. We found that when the length of VMFs was 5 aa, numerous HFs were obtained for many VMFs. When the length of VMFs was extended to 7 aa, approximately 97% of queries led to identical subjects. When the length was further extended to 9 aa or 11 aa, less than 7% of the queries led to matches. Notably, regardless of whether the length was 7 aa or more aa, the main conclusion of this study would not be changed. Therefore, VMFs containing 7 aa (XXXMXXX) were selected for the original analysis database creation, with “M” indicating one mutation and “X” indicating the adjacent aa (Table S2). Among the subjects obtained, only those fulfilling all of the following criteria were collected: a) 100% VMF identity, that is, the 7 aa was fully recognized in the database; b) 100% homologous sequences; and c) sequence origins from SARS proteins were ignored. At this stage, we obtained approximately 8000 subjects in total. To prevent duplicative statistics and overcome the limitation of the lower error, we defined two filter conditions: a) for multiple accessions for one protein, only one accession was retained, and b) for multiple isoforms for same one protein, only one isoform was retained. After filtering, 5600 subjects were collected and were used to establish the final database for analysis, and it included the name, ID number, species, and submission number for each HF (Fig. 1c, Tables S3-1, S3-2, S3-3, S4).

### Bacteria contributed more than 60% of the homologous fragments

The species proportion in the final database was analyzed. We were astonished to find that bacteria contributed more than 60% of the HFs, while Animalia, Fungi, Plantae and Protista accounted for 19, 11, 4 and 2%, respectively (Fig. 1d). Notably, of the 742923 subjects in the NCBI protein database, more than 93.5% were found in eukaryotes, and fewer than 5% were found in bacteria (Fig. 1e, Fig. S2). The markedly inverse proportion inspired us to pay close attention to the bacteria. The species proportions were further analyzed based on individual mutation location, and no significant difference was found among S, E and M protein mutations (Fig. 1d). Then, the database was analyzed from the perspective of individual mutations. We found that the number of HFs ranged from 0 to 2075, averaging 92 for each of the 61 evaluated mutations. No HFs were found for 4 mutations (T716I, D80A, G142D and G339D), accounting for only 7% of the total evaluated mutations. Interestingly, although bacterial HFs were identified for 93% of the mutations, for each mutation, the bacterial proportion was not correlated with the number of HFs. For certain mutations, both the identified HF and bacterial contributions were very high; these mutations included S447N (209), T20N (119), T76I (264), and P71L (1523). In contrast, fewer than 5 HFs were found for some mutations, such as N501Y (5), L452R (3), and T716I (1), but all of the HFs were from bacteria. Notably, the bacterial proportion for N679K was is 0%, and for the other 6 mutations, although the corresponding HF number was high, it was 13, arguing against the idea that the bacterial contribution that was revealed was a coincidence (Fig. 1f).

### The human microbiota is a reservoir of homologous fragments

Next, we tried to understand the composition of the bacteria obtained via categorization and performing a quantitative analysis. Based on phylogeny and taxonomy theories, the 3271 obtained bacterial species belonged to 73 different phyla. Despite the diversity of these phyla, most of the bacterial species obtained were classified into four major bacterial phyla, namely, Proteobacteria, 41%; Actinobacteria, 29%; Bacteroidetes, 8% and Firmicutes, 8% (Fig. 2a). Notably, the four major phyla containing HFs were consistent with the four major phyla of human microbiota, although the proportions were different ^38,39^. The composition of the bacterial species with S, M, and E protein mutations was then analyzed individually, and a composition pattern similar to that of the mutations in total was obtained. The correlations between the mutations and bacterial phyla with proportions greater than 1% were plotted. We found that all 61 mutations matched multiple bacterial phyla (Fig. 2b). For example, HFs carrying I82T were found in descending order in Proteobacteria, Bacteroidetes, Firmicutes and Actinobacteria, while HFs containing T9I were found in descending order in Bacteroidetes, Actinobacteria and Firmicutes. In addition, some bacterial phyla carried multiple HFs, including Proteobacteria, which contributed more than 51 different types of HFs (Fig. 2b). To deepen our understanding of the composition of the contributing bacterial, core species were further categorized at the family and genus levels. The four major bacterial phyla contained 331 families, including 132 in Proteobacteria, 89 in Actinobacteria, 30 in Firmicutes and 28 in Bacteroidetes (Fig. S3a). At the genus level, Streptomyces, and genera in the Pseudomonadaceae, Enterobacteriaceae, Micromonospora, Xanthomonadaceae, Rhizobiaceae, Burkholderiaceae, Flavobacteriaceae, and Comamonadaceae families, showed average HF abundances of ≥ 1%, and therefore, they were considered to be the top ten HF-contributing genera (Fig. S3b).

**Fig. 2.**
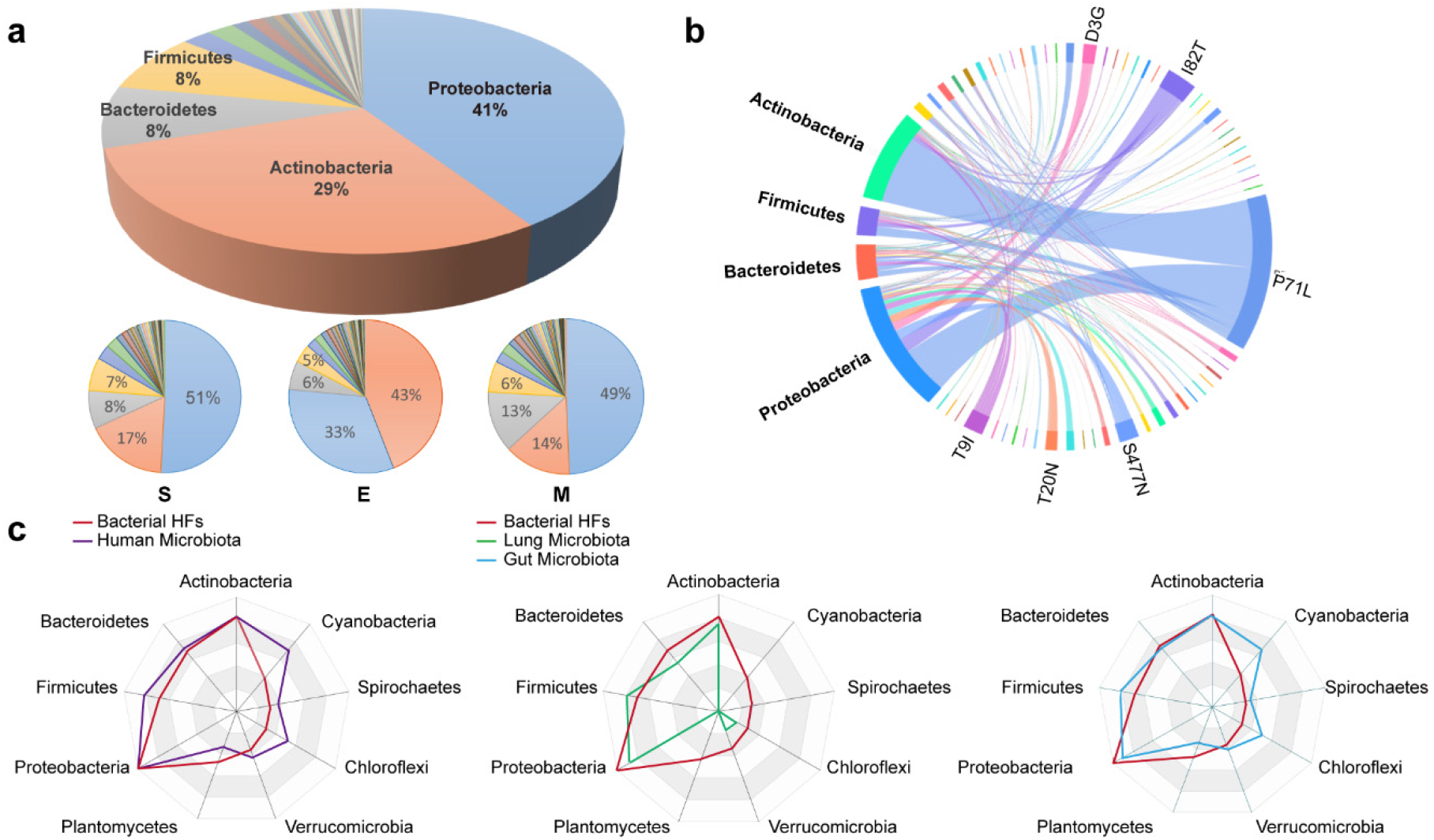
Homologous fragment (HF) composition analysis of the obtained bacteria. **a,** Pie chart representations of the obtained bacteria, all items (up) and items of spike, envelope and membrane proteins (bottom), at the phylum level. **b**, Chord diagram displaying the network of mutated HFs and their bacterial phyla. **c,** Radar maps showing the correlation of the identified bacteria sources of HFs (red), human microbiota (purple) (top), lung microbiota (green) (below left), and gut microbiota (cyan) (below right) at the phylum level.

As the largest host of SARS-CoV-2, the human body harbors trillions of microbial species that colonize the skin, oral cavity, intestines and even the lungs ^40–43^. The four main phyla between the human microbiota and the bacteria were identical, enabling deep correlation analysis. As shown in the radar chart, almost all bacteria were belonged to the human microbiota at the phylum level. In COVID-19 patients, the lung is a major affected organs due to viral infection or viral-bacterial coinfection. Approximately 56% of the bacteria we obtained are found in the lungs of healthy individuals; these include Streptomycetaceae and Staphylococcaceae (Fig. 2c)^40,44^. Although bacteria in lung are abundant, microbial cell populations reach their highest density in the intestines, where they form the gut microbiota and are believed to be important to human life ^41,42^. Interestingly, 83% of the obtained bacterial species were also found in the human gut microbiota database (Fig. 2c), implying the importance of the gut microbiota.

### The species composition of the homologous fragments is similar to that of the human gut microbiota in COVID-19 patients

In addition to pneumonia, diarrhea and gastrointestinal symptoms were found in 10% of COVID-19 patients ^45,46^. It has been reported that in approximately 50% of COVID- 19 patients, the virus was found in feces, leading to the hypothesis that there is not only replication and therefore activity in the intestine but also that the virus resides for prolonged periods in the intestines ^47–49^. Increasing evidence suggests that gut bacteria are altered in COVID-19 patients, and these alterations are associated with infection severity, treatment effectiveness, and prognosis ^50,51^. The gut microbial ecological network is significantly weakened and becomes diffuse in patients with COVID-19, and together, the number of beneficial bacteria is decreased, and harmful bacteria production multiplies ^50,52–54^. Two gut microbe databases of COVID-19 patients (gutMEGA) were analyzed. Compared with the composition of non-COVID-19 gut microbes, two abnormal changes were detected in the COVID-19 entries. The population of beneficial bacteria, particularly Firmicutes, significantly had declined to 47% from 72%, while that of harmful bacteria, such as Proteobacteria, increased to 9% from 1%. Notably, a similar change was observed for the obtained bacteria carrying HFs, including an increase in harmful Proteobacteria to 41% and a reduction in beneficial Firmicutes to 8% (Fig. 3a) ^50,51,54^. The influences of gut bacteria changes identified in the two databases were perhaps best exemplified by an intersection analysis with the bacteria containing S protein mutations (Fig. 3b). In the two COVID- 19 patient databases, at the family level, 17 and 11 coexpressed bacteria were identified; at the species level, 3 coexpressed bacteria were identified. These correlations at the genus and species levels were further analyzed (Fig. 3c). At the genus level, we found that among the 55 S mutations, 15 were detected in 19 differential bacterial genera in COVID-19 patients. At the species level, 3 types of bacteria in COVID-19 patients were found to carry mutational HFs (Fig. 3c). Notably, one genus of bacteria, Escherichia, carried 3 different mutations (S371P, S373P and S477N). Considering the limited size of the current gut bacteria database of COVID-19 patient data, these high correlations reinforced the idea that SARS-CoV-2 may acquire HFs from the human microbiota.

**Fig. 3.**
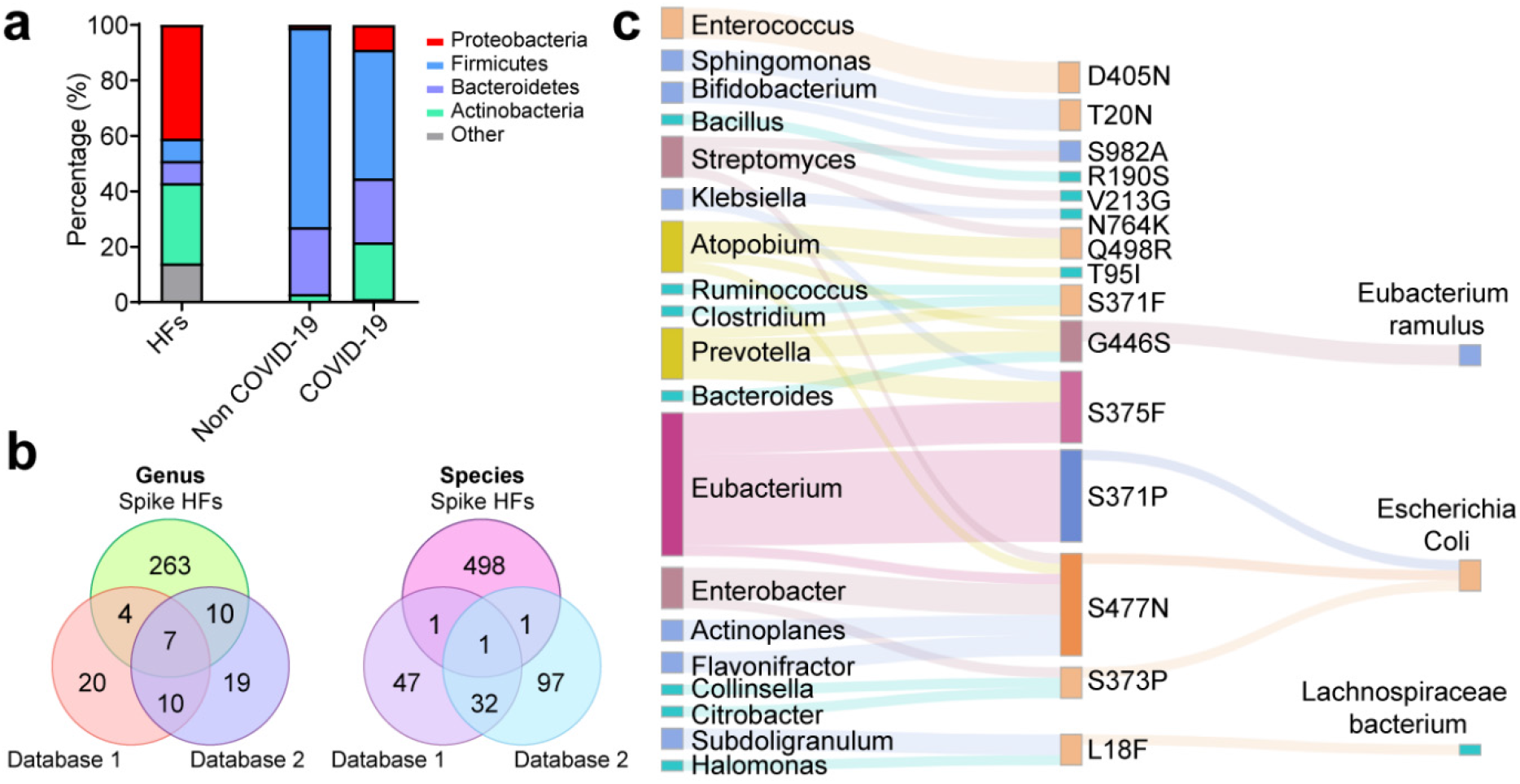
Correlation of the identified bacteria carrying homologous fragments and those in the human gut microbiota in COVID-19 patients. **a,** Abundances of microbial phyla detected in HFs and stools from in-hospital patients with COVID-19 and non-COVID-19 individuals. The average relative abundances of COVID-19 and non-COVID-19 individuals were obtained from two databases (gutMEGA). **b,** Intersection analysis showing the obtained bacteria containing HFs with the gut microbes of COVID-19 patient databases at the genus (left) and species (right) levels. Two databases were used (gutMEGA). **c,** A Sankey diagram showing the relationship of the mutations in the spike protein with changes in bacteria containing HFs in COVID-19 patients at the genus (left side) and species (right side) levels.

After the emergence BA.5, various new omicron subvariants appeared, including BA.5.2 and BF.7, two major pandemic variants, and BQ.1.1 and XBB, the most resistant SARS-CoV-2 variants discovered to date. These new recombinant subvariants carried 5 new advantageous mutations, as determined on the basis of the original Omicron, with mutation frequencies higher than 90%, including R346T, K444T, N460K, and F486S in the S protein and T11A in the E protein (Fig. S4a). We applied our VMF-based blast model to search for potential HFs carrying these five mutantions. As expected, bacteria contributed more than 80% of the HFs with these five mutations, while Animalia, fungi, Plantae and Protista accounted for 4, 4, 5 and 1%, respectively (Fig. S4b, Table S5). Moreover, the identified bacterial species were also classified into four major bacterial phyla, namely, Firmicutes, 49%; Proteobacteria, 26%; Actinobacteria, 11%; and Bacteroidetes, 4% (Fig. S4c), which were the same as those carrying mutations previously identified.

### SARS-CoV-2 RdRp replicates bacterial mRNAs to form chimeric viral-bacterial RNAs harboring mutations

In contrast to SARS-CoV-2, bacterial HFs are encoded by mRNAs. Nucleotide sequences of bacterial mRNAs encoding exactly the same amino acid sequences as viral RNAs were aligned, revealing a percent identity (PNI) from 63% to 90% in the 7 examined fragments (Fig. S5). A PNI less than 100% can be possibly due to the degeneracy of the genetic code and particularly the presence of synonymous mutations in SARS-CoV-2 (GenBank accession: NC_045512.2; Wuhan-Hu-1) ^55,56^. SARS-CoV- 2 replication is mediated by a multisubunit replication-and-transcription complex of viral nonstructural proteins (nsp), of which nsp12 is the core component of the RNA- dependent RNA polymerase (RdRp) ^3^. When these bacterial mRNAs introduce mutations into a viral genome, they must serve as templates for the viral RdRp complex, which was examined experimentally (Fig. 4a). The bacterial mRNAs (up to 27 nt long) carrying mutations such as N501Y, E484K and L452R, which were fused with a previously validated viral duplex RNA, were successfully replicated by an RdRp complex containing nsp12, nsp7 and nsp8 (Fig. 4b, Table S6). This experiment demonstrated that exogenous bacterial mRNAs are effective templates for SARS-CoV- 2 RdRp.

**Fig. 4.**
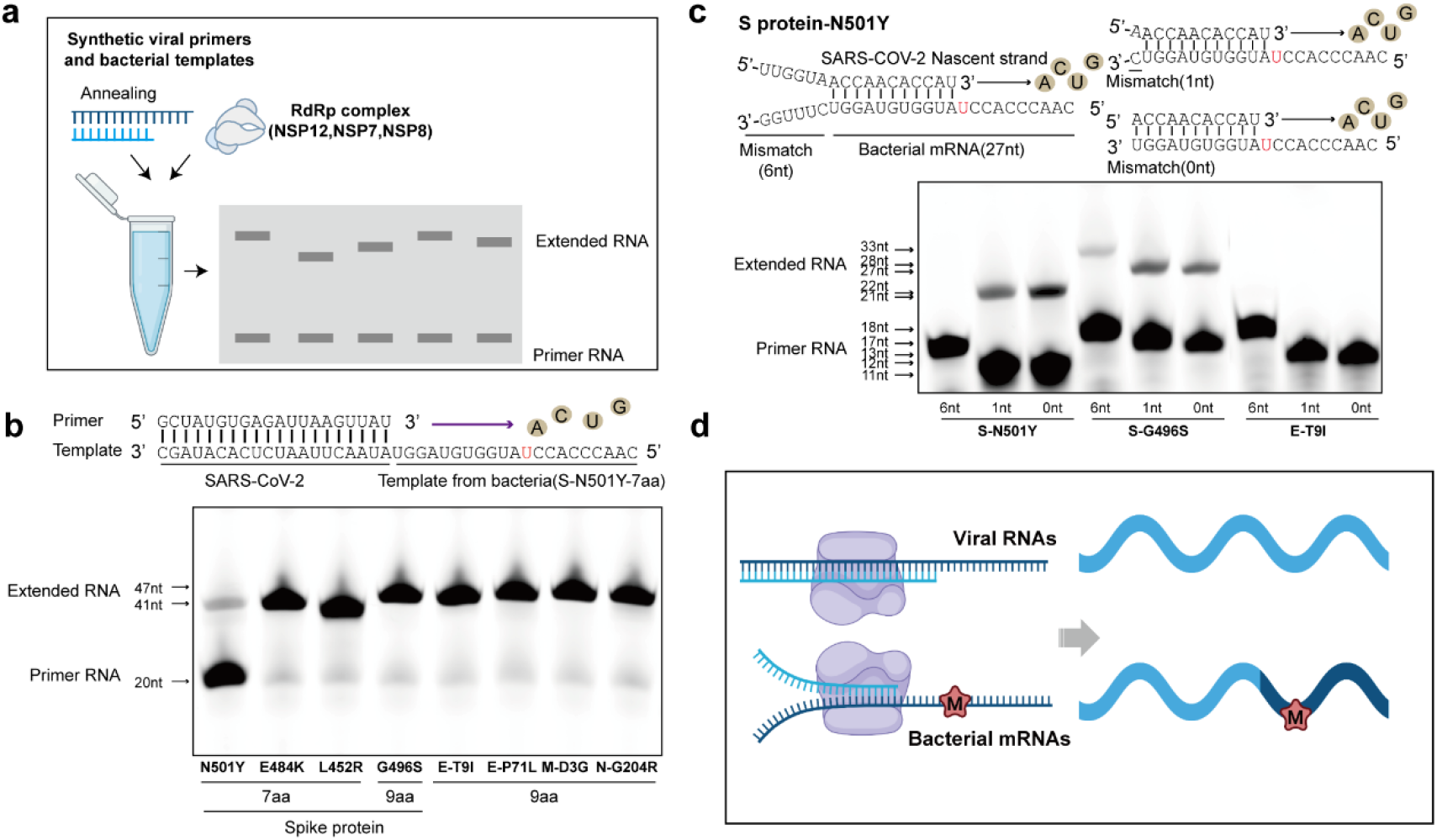
SARS-CoV-2 RdRp introduces bacterial mRNA nucleotide mutations into viral RNAs through homologous recombination. **a,** Schematic diagram showing the RdRp extension experiment. **b,** RNA duplex with a 5′-21 nt or 27 nt from bacterial mRNA overhang as the template for primer extension and RdRp-RNA complex assembly (exemplified by N501Y). The primer strand is labeled with a fluorescence molecule at the 5’ end. **c,** The primer strand from SARS-CoV-2 contains 11 or 12 nucleotides that complementarily pair with bacterial template mRNAs but have 0, 1 or 6 mismatches at the 5’ end and in a duplex formation with the bacterial template mRNAs harboring N501Y, G496S or T9I mutations for RdRp extension (exemplified by N501Y). **d,** Schematic diagram showing RdRp introducing a nucleotide mutation into a bacterial mRNA to produce a bacterial-viral chimeric RNA via homologous recombination.

During SARS-CoV-2 replication, base pairings with 6-12 consecutive nucleotides can serve as “junction” sites for template switching, i.e., RdRp switches from copying one genome to copying another, resulting in a subgenome ^57–60^. Notably, under certain circumstances, such as when lesions are included in a template, replication-impeding secondary structures are formed, and the tightly bound nucleotide pool is imbalanced, template switching may also be evident among RNAs of different species ^59,61,62^. Whether the complementary base pairs between bacterial mRNA and viral RNA can act as “junction” sites was therefore examined. As shown in Figure 4c, the nascent primer strand from SARS-CoV-2 contained 11/12/17 nucleotides; three bacterial template mRNAs harboring the N501Y mutation carried 11 nucleotides complementary to the viral primer but 0, 1 or 6 mismatches in the 5’ end. Among the three nucleotide groups, only the last group failed to extend. Then, two more types of bacterial template mRNAs harboring G496S (12 complementary nt) or T9I (12 complementary nt) were individually examined under identical conditions. Extension was observed for all G496S mRNAs regardless of the number of mismatches in the 5’ end but extension failed for all the T9I mRNAs (Fig. 4b-d, Table S6). The different replication efficiency was perhaps due to the insufficient efficacy of RdRp in the *in vitro* system. Although these results demonstrated that nucleotide mutations in bacterial mRNAs can be introduced into viral RNAs during replication via the RdRp, clear evidence showing that mutations can be integrated into the SARS-CoV-2 genome was not found. Because one cannot rule out the possibility that artificial coinfection of live virus and bacteria in laboratories may promote the emergence of new mutations with unpredictable capabilities, experiments with viral RNA and live bacteria is currently prohibited at our organization.

Since Dec. 2019, the number of infections has reached 700 million worldwide. The ultrahigh number of infections and wide regions of infection may have provided a wealth of bacterial carriers for the omicron variant that was not available for other variants. Although data errors cannot be ruled out due to different infection density data and variable persistence of gene surveillance for different variants, all the evidence provided above supports the idea that SARS-CoV-2 may acquire HFs from the human microbiota, as is summarized in a schematic graph (Fig. 5). In comparison with the original strain, SARS-CoV-2 VOCs have evolved into chimeras containing components from the human microbiota.

**Fig. 5.**
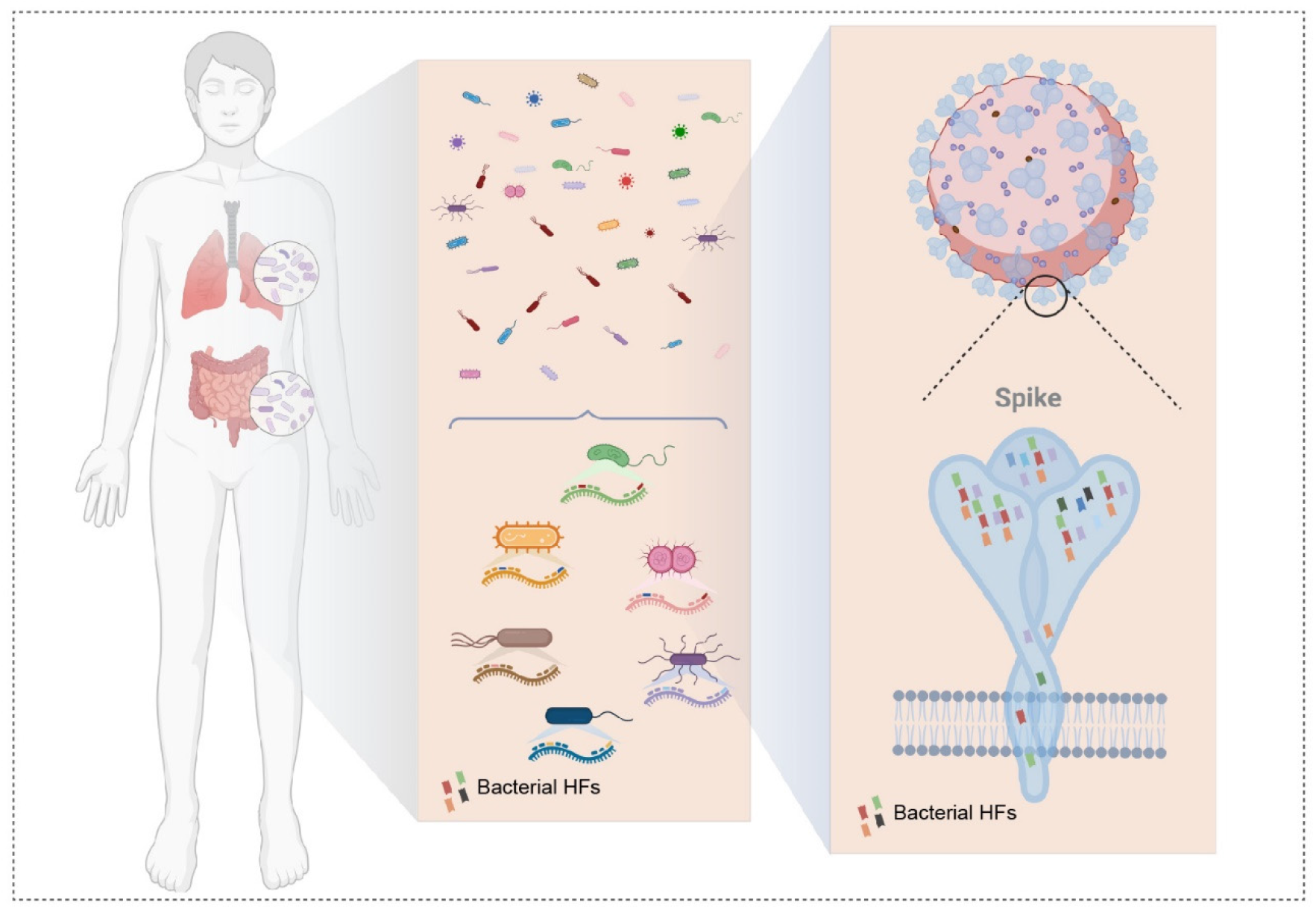
SARS-CoV-2 VOCs are chimeras containing homologous fragments (HFs) from the human microbiota. Schematic graph showing SARS-CoV-2 acquiring HFs from human microbiota as exemplified by the S protein.

## Discussion

Available evidence supports the idea that SARS-CoV-2 shows high capacity to acquire advantageous mutations ^6,18^. Here, a VMF-based blast analysis was performed to explore sources of high-frequency advantageous mutations identified in VOCs. Among the identified species containing HFs, more than 70% were derived from the human microbiota. In particular, the changes in the pattern of gut bacteria in COVID-19 patients are quite similar to those of the bacteria we identified harboring HFs. In addition, nucleotide mutations in bacterial mRNAs can be introduced into viral RNAs during replication via RdRp. We propose that the human microbiota is the potential template reservoir involved in SARS-CoV-2 mutation accumulation.

In the possible case of SARS-CoV-2 acquiring mutations from human bacteria, at least three possible scenarios explain viral access to bacterial mRNA. In Scenario One, exposure of bacteria and virus are both exposed to phagocytes, including macrophages, neutrophils, etc. Bacterial coinfection is a common complication of SARS-CoV-2 infections. In particular, severely affected patients suffer with a significantly higher rate of coinfection with bacteria (26%) ^63^. The coinfecting bacteria include mainly *Acinetobacter baumannii*, *Klebsiella pneumoniae*, and *Pseudomonas aeruginosa*, which were also identified in our study. During pathogen invasion, macrophages are among the first immune cells to respond. As scavengers of the immune system, they show the profound ability to engulf microorganisms and digest them. Therefore, macrophages are probable hosts for both bacterial and viral genomes during coinfection ^64–66^. In Scenario Two, bacterial mRNA is exposed to nonphagocyte cells, such as infected cells or impaired cells. During SARS-CoV-2 infection, the immune barrier is impaired, which may cause invasive pathogens such as Streptococcus to break through the immune barrier and invade nonphagocytic cells ^67,68^. The invaded host cells can target intracellular bacteria through autophagic machinery to lyse the bacteria and block bacterial proliferation. However, autophagy that fails to control the replication and spread of bacteria may trigger phagocytosis by phagocytes. At this point, the bacterial genome and the viral genome may coexist in host cells or phagocytes. In addition, the genetic information from the lysed bacteria may be engulfed by infected cells in some cases ^69–71^. In Scenario Three, some bacteria might be potential hosts for the virus. The mechanism underlying phage (a group of viruses that infect bacteria) invasion of bacteria has been elucidated and has become a reliable gene-editing method ^72^. In addition, certain bacterial strains can increase the rate of viral coinfection in mammalian cells, facilitating genetic recombination between two different viruses and thus removing deleterious mutations and restoring viral fitness ^73^. Recently, studies have shown that some ciliates, a type of single-cell organism, consumed virus particles, which fostered their population growth, revealing an unexpected role for viruses in the ecosystems ^74^. As the two most abundant microbial entities, bacteria and viruses coexistence remains puzzling. These scenarios provide infinite possibilities for virus– bacteria interactions during the mutational evolution of SARS-CoV-2.

SARS-CoV-2 reference genome analyses have revealed multiple sudden changes in viral sequence during evolution, signaling potential recombination events ^6,18,21^. The present study supports the idea that the complementary base pairs between bacterial and viral RNAs promote the recombination and production of chimeric RNAs harboring mutations. For RNA viruses, homologous recombination is the most common type of recombination ^75,76^. RNA‒RNA interactions are the determinants for the template switching efficacy among SARS-CoV-2 RNAs, with 6–12 consecutive complementary nucleotides serving as “junction” sites. Notably, the longest bacterial HF we identified was 9 aa, indicating that as many 27 consecutive complementary nucleotides can be shared between bacterial and viral RNAs. In a less-than-ideal *in vitro* experimental system used in the present study, we showed that 11 consecutive complementary nucleotides were sufficient to generate chimeric RNAs resulting from template switching. During homologous recombination, nucleotide mutations were simultaneously introduced into the viral chimeric RNA. Notably, not all mutations were matched to an HF (Fig. 1f). Therefore, although our study emphasized the importance of RdRp-mediated homologous recombination between bacterial and viral RNAs, the results are also compatible with the replication-associated mutation and evolutionary pressure selection theories ^55,77–79^.

COVID-19 is rapidly spreading worldwide. Scientists are monitoring recombination and transmission events in the SARS-CoV-2 population in real time and exploring the evolution of SARS-CoV-2 from its initial introduction into human populations to the present day and have been exploring the mechanisms underlying SARS-CoV-2 variation acquisition. Our research broadens the perspective to include virus and microbiota studies but leaves many questions unanswered, such as how and where does SARS-CoV-2 access bacteria *in vivo*, and how are chimeric RNAs harboring mutations integrated into the SARS-CoV-2 genome? Whether bacteria can be potential hosts in the process of virus transmission, and the relationship between viruses, bacteria, and hosts deserves further exploration. Our findings are based on the analysis of acquired mutations in the virus, which was insufficient to explain bidirectional causality between viruses and bacteria. In addition, due to multiple factors and experimental restrictions, we were unable to provide more experimental evidence, but we hope that more researchers in this field can perform in-depth investigations. Our work shows enormous implications for comprehending the future trajectory of the COVID-19 pandemic and developing preventive and treatment strategies.

## Acknowledgement

We thank Prof. Yangbo Hu (Wuhan Institute of Virology, CAS) and his team for the exploration on engeenered bacteria experiments. We are grateful to the National Science Fund of Distinguished Young Scholars (81825021), Fund of Youth Innovation Promotion Association (2019285), the National Natural Science Foundation of China (31700732, 81773707, 92169202), the National Key Research and Development Program of China (2020YFC0842000), the National Key Laboratory Program of China (LG202101-01-04), Fund of Shanghai Science and Technology (20ZR1474200, 22QA1411000).

## Author contributions

Z. G., B.X., and J.L. conceived the project. Z.G. and B.X. designed the experiments; B.X. and B.C. carried out bioinformatic analysis. B.C. performed the RdRp extention experiments; W.Y. and H.X. purified the proteins; all authors analyzed and discussed the data. Z.G., B.X. and B.C. wrote the manuscript. All authors read and approved the manuscript.

## Methods

### Identification of HFs

Homologous fragments (HFs) identification was carried out in NCBI Protein database. VMFs (7aa) was submit to NCBI protein blast research website. The alignment parameters were set as Database Non-redundant protein sequences (nr), Max target sequences 5000, Expect threshold 0.05, Word size 7. Among the obtained subjects, only those fulfill all the following criteria were collected: a) 100% VMF identity, that is the query 7 aa could be fully recognized by the database; b) 100% homologous sequences; c) sequence origins from SARS proteins were ignored. In order to avoid duplicate statistics and overcome the limitation of the lower error, we further defined two filter conditions: a) multiple accessions for one protein, only one accession was retained; b) multiple isoforms for same one protein, only one isoform was retained. After filtering, the name, ID number, species, submission number were collected.

### Sequence data processing, inferring gut microbiota composition and statistical analysis

The composition of the obtained HFs bacteria through categorization and quantitative analysis through NCBI database and Uniprot database. Following this, microbiota composition profiles were inferred from quality-filtered forward reads using MetaPhlAn218 V.2.7.7 with the V.20 database. Associations of HFs microbial species with human microbiota parameters were identified using gutMEGA, GMrepo, China national Genebank Microbiome, HOMD Human oral microbiome Database V3, NIH Human Microbiome Project.

### RNA extension assays

All RNA oligonucleotides (templates and primer) were chemically synthesized by Genscript, in which the primer had a FAM-labelled at the 5’ end. Template and Primer oligonucleotides were annealed at 1:1 molar ratio by a 5min heating at 85°C followed by gradually cooling to room temperature in the annealing buffer (10 mM Tris-HCl, pH 8.0, 2.5 mM EDTA, and 25 mM NaCl). The primer sequences were listed in Table S6. RNA extension reactions contained annealed RNA (2.7 μM), nsp12 (2.7 μM), nsp7 (6.75 μM) and nsp8 (6.75 μM) in reaction buffer (20 mM Tris, pH8.0, 10 mM KCl, 6mM MgCl2, 0.01% Triton-X100, and 1mM DTT) with 10 mM NTP (2.5mM ATP, 2.5mM UTP, 2.5mM GTP, 2.5mM CTP) and 1.14 U/ul RNA inhibitor. The total volume of the reaction solution was 20μl. After incubating at 25°C for 1.5 hours, 40μl quench buffer (94% formamide, 30 mM EDTA, prepared with DEPC-treat water) was added to stop the extension reaction. Sample was prepared with 8μl reaction solution and 2μl 5×DNA loading, then loaded onto a 15% Urea-PAGE denatured gel, run at 90V for 1.5h, and visualized by Typhoon 95000 FLA Imager.

## Supplemental Figures and Table

**Table S1. Frequency of all mutations in 5 VOC variants.** All mutations have frequencies above 90%, which were referred to CNCB-NGDC RCoV19, NCBI Coronavirus Resource and EBI coronavirus database.

**Table S2. The HFs sequences of spike, envelop and membranes proteins. 7**aa HFs were composed of mutation and its adjacent 6 amino acids (aa).

**Table S3-1. Original data for HFs on spike protein.**

**Table S3-2. Original data for HFs on envelope protein.**

**Table S3-3. Original data for HFs on membrane protein.**

**Table S4. Original data for 9aa HFs on spike, envelope and membranes proteins.**

**Table S5. Original data for HFs on Omicron subvariants, including BA.5.2, BF.7, BQ.1.1 and XBB.**

**Table S6. The primer list for RdRp extension assays.**

**Fig.S1.**
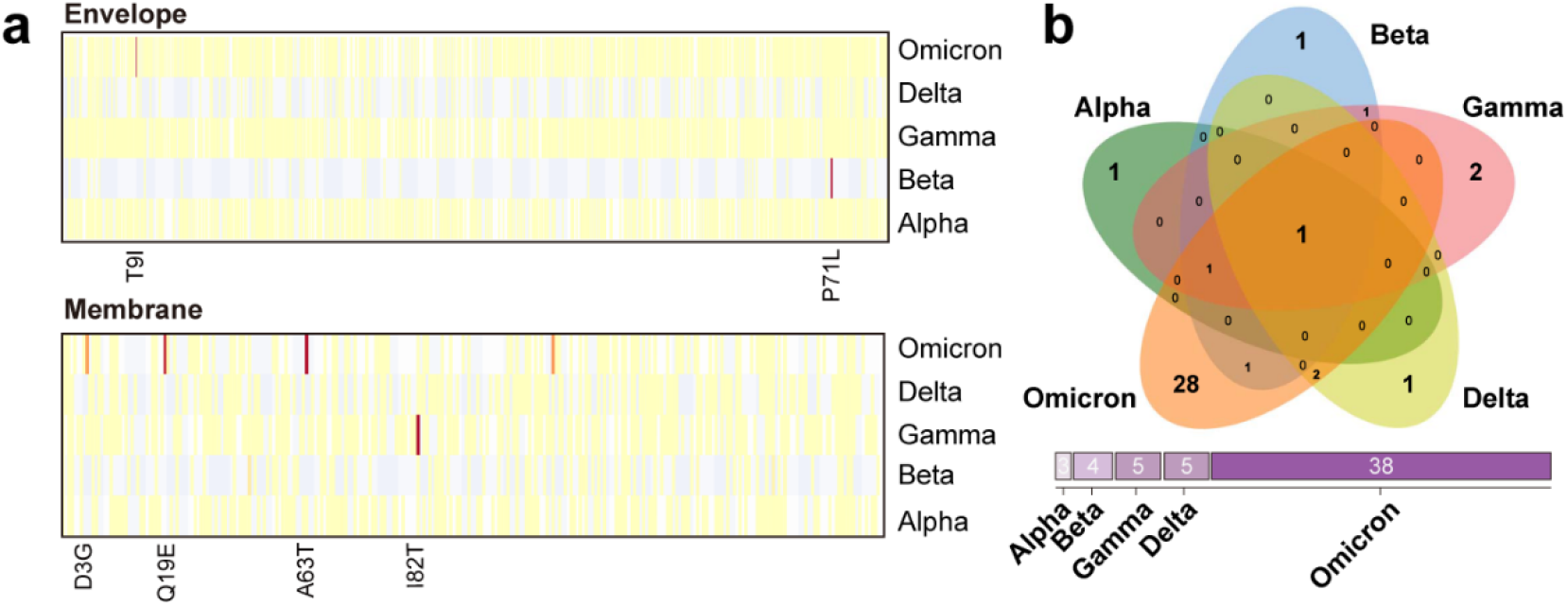
Advantageous mutaions of VOCs. **a,** The mutation probability of envelope and membrane protein mutations in all VOCs. **b,** Comparison of high frequent mutations in different VOCs.

**Fig.S2.**
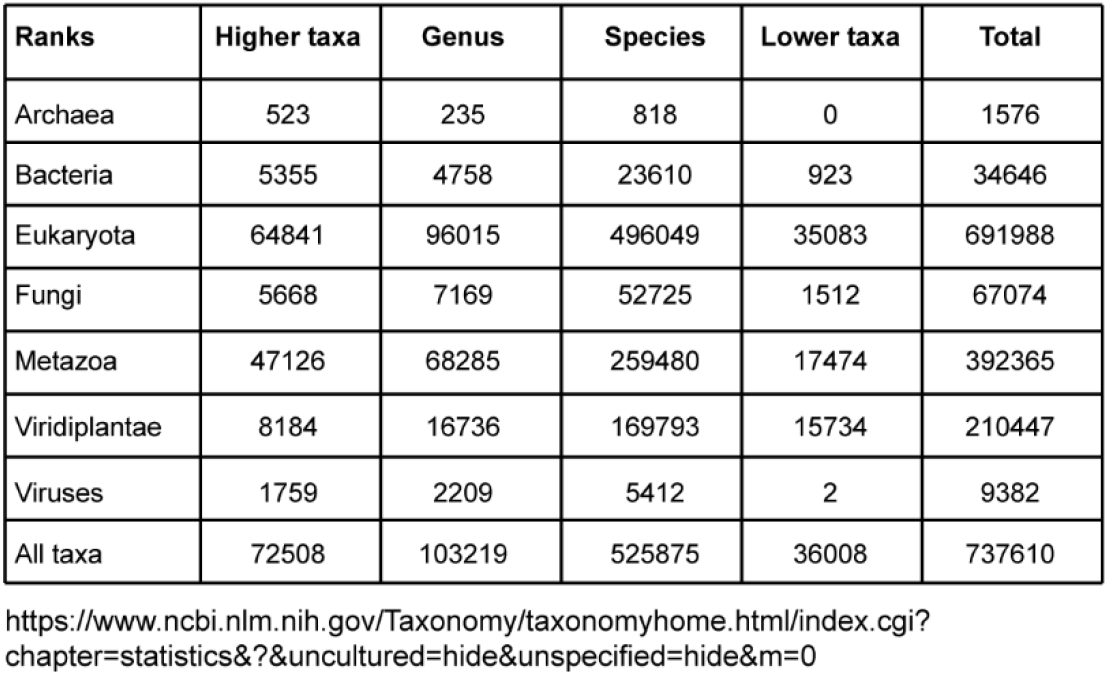
Species taxonomy in NCBI database. The database currently represents about 10% described species of life on the planet (October 2022).

**Fig.S3.**
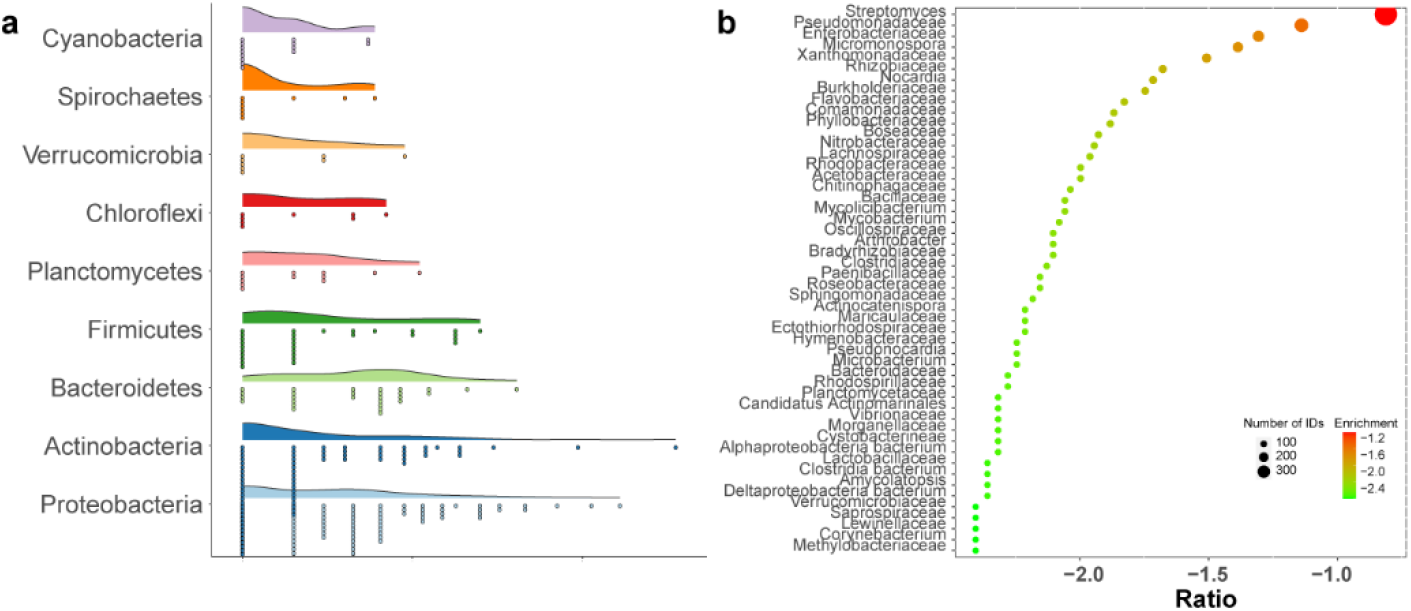
The HFs composition analysis of obtained bacteria at family and Genus levels. **a,** Composition of top bacteria phylum at family level. The raindrops represent the family types of each phylum. **b,** Enrichment analysis of the obtained bacteria at family level. Color coded with log10 transformed of family percentage.

**Fig.S4.**
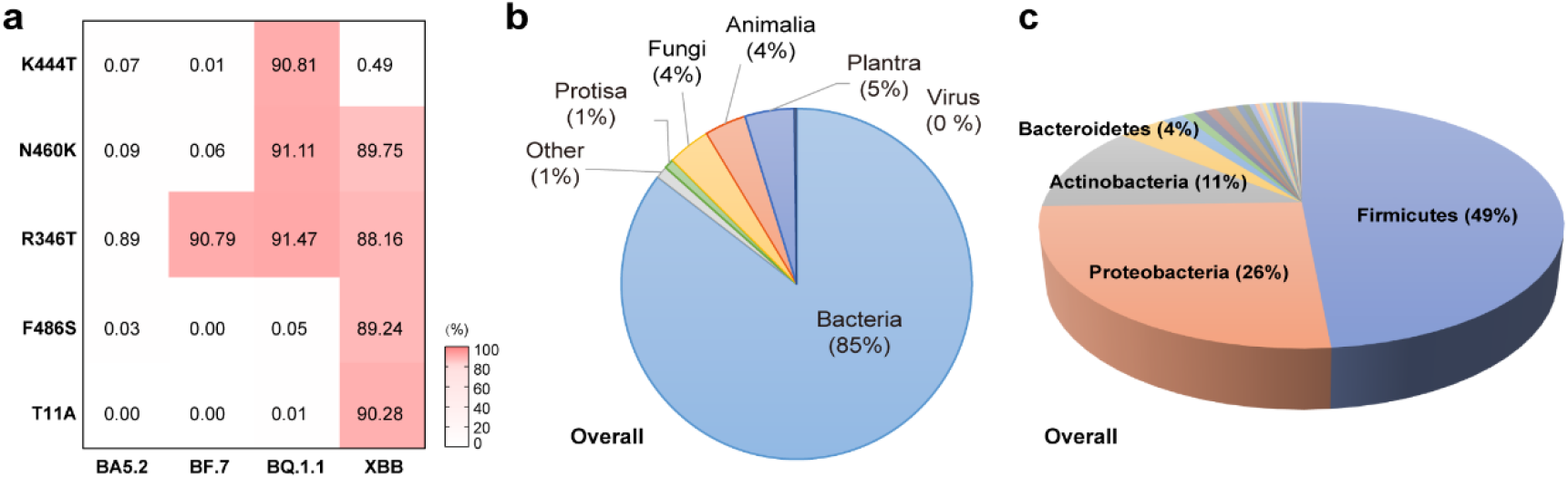
Homologous fragments (HFs) alignment of new mutations fragments on the Omicron subvariants. **a,** Heatmap of 5 advantageous mutations frequencies in Omicron subvariants. **b,** Pie chart representations of HFs Kingdoms from overall (bigger one), and four mutations on spike (S), one mutation on envelope (E) (smaller five). The species proportions in HFs and NCBI databases. **c,** Pie chart representations of the obtained bacteria, all items at phylum level.

**Fig.S5.**
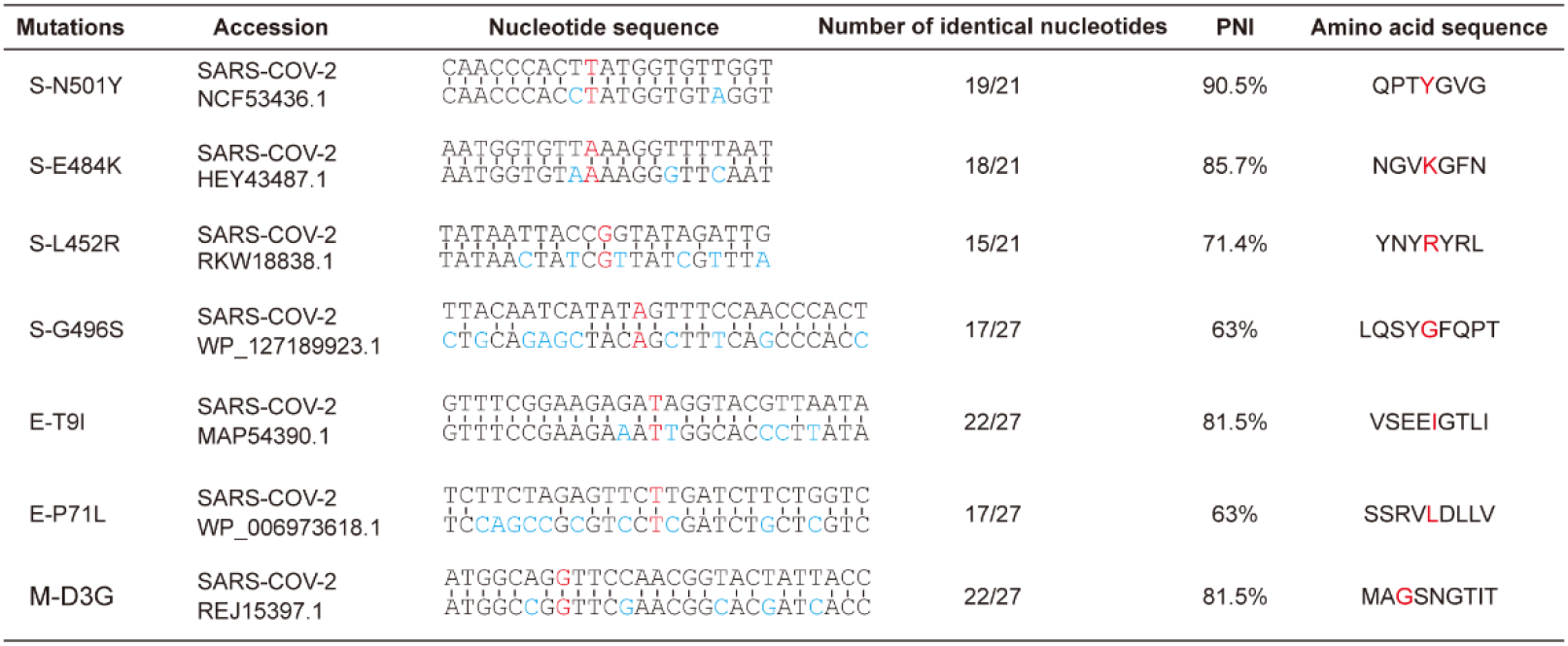
The alignment of nucleotide sequences of bacterial mRNAs that encoding same amino acid sequences with viral RNAs, including the mutations from S protein (N501Y, E484K, L452R and G496S), E protein (T9I,P71L) and M protein (D3G). Red represents mutated base or amino acid, blue represents inconsistent bases. PNI, The percent nucleotide identity.

